# Oral Fentanyl Consumption and Withdrawal Impairs Fear Extinction Learning and Enhances Basolateral Amygdala Principal Neuron Excitatory-Inhibitory Balance in Male and Female Mice

**DOI:** 10.1101/2023.11.28.569085

**Authors:** Anthony M. Downs, Gracianne Kmiec, Zoé A. McElligott

## Abstract

The number of opioid overdose deaths has increased over the past several years, mainly driven by an increase in the availability of highly potent synthetic opioids, like fentanyl, in the un-regulated drug supply. Over the last few years, changes in the drug supply, and in particular the availability of counterfeit pills containing fentanyl, have made oral use of opioids a more common route of administration. Here, we used a drinking in the dark (DiD) paradigm to model oral fentanyl self-administration using increasing fentanyl concentrations in male and female mice over 5 weeks. Fentanyl consumption peaked in both female and male mice at the 30 µg/mL dose, with female mice consuming significantly more fentanyl than male mice. Mice consumed sufficient fentanyl such that withdrawal was precipitated with naloxone, with males having more withdrawal symptoms, despite lower pharmacological exposure. We also performed behavioral assays to measure avoidance behavior and reward-seeking during fentanyl abstinence. Female mice displayed reduced avoidance behaviors in the open field assay, whereas male mice showed increased avoidance in the light/dark box assay. Female mice also exhibited increased reward-seeking in the sucrose preference test. Fentanyl-consuming mice of both sexes showed impaired cued fear extinction learning following fear conditioning and increased excitatory synaptic drive and increased excitability of BLA principal neurons. Our experiments demonstrate that long-term oral fentanyl consumption results in wide-ranging physiological and behavioral disruptions. This model could be useful to further study fentanyl withdrawal syndrome and behaviors and neuroplasticity associated with protracted fentanyl withdrawal.

## Introduction

Opioid Use Disorder (OUD) has become increasingly prevalent over the last decade with over 26 million people worldwide affected by OUD and 110,360 overdose deaths in March 2022 in the US alone ^1,2^. These overdose deaths were primarily attributed to changes in the drug supply and increased availability of highly potent synthetic opioids, such as fentanyl and its analogs ^3^. OUD is a chronically relapsing condition characterized by cycles of craving, binge, and withdrawal ^4^. In OUD, excessive opioid use occurs, in part, as a negative reinforcer, to prevent the physical and affective components of withdrawal. Due to recent changes in the drug supply, it is especially important to develop new models to explore withdrawal syndromes associated with fentanyl use to help facilitate the development of new treatments for this disorder.

In the present study, we developed a drinking in the dark (DiD) mouse model of oral fentanyl consumption. While many animal self-administration studies have used intravenous self-administration, it is important to consider that many people who use opioids use other routes of administration. Oral routes of administration are increasingly common with opioids, especially early in the history of use ^5^. This may be an especially important consideration for fentanyl, where many users self-report a preference for smoking or oral consumption as a route of administration ^6–8^. Ingestion and smoking fentanyl have become even more common with the prevalence of fake pills containing fentanyl that have flooded the market ^9,10^. To date, there have been a number of animal studies that have demonstrated oral consumption of opioids in rodents using both operant and home-cage drinking resulting in neural and behavioral adaptation ^11–24^.

Previous studies in our laboratory have utilized a model of precipitated morphine withdrawal to investigate the somatic and negative affective aspects of opioid withdrawal. Notably, we have demonstrated significant sex-dependent effects on both neuroplasticity and behavior in response to opioid withdrawal ^25–27^. Previous work from other laboratories have identified fear extinction learning deficits following opioid administration, which may be important to understand well-established links between Post Traumatic Stress Disorder (PTSD) and opioid use disorder ^28–33^. In the present study, we expand on prior research by developing a single-bottle home-cage drinking-in-the-dark (DiD) model of oral fentanyl consumption. We investigated fentanyl consumption across a range of concentrations, somatic withdrawal signs, behavioral responses in approach/avoidance assays, fear learning and extinction, and basolateral amygdala (BLA) physiology during fentanyl consumption in both male and female mice.

## Methods

### Subjects

All Procedures were approved by the University of North Carolina Institutional Animal Use and Care Committee. Male (n=24) and female (n=24) C57BL/6J mice (The Jackson Laboratory, Bar Harbor, ME) at 8 weeks in age were used in all experiments and singly housed. All animals had food and water *ad libitum* for the duration of the study. After the completion of the study, all animals were euthanized according to IACUC protocols.

### Experimental Design

Experiments were conducted with 4 cohorts of 16 mice each, with each cohort evenly split between water and fentanyl groups and males and females. Cohorts 1, 2, and 3 were habituated to the reverse light cycle for 1 week and then underwent 5 weeks of fentanyl drinking. Mice then underwent behavioral testing beginning after 3 days of fentanyl abstinence in the following order: open field, elevated plus-maze, light-dark box, sucrose-preference test, and fear conditioning. Cohort 4 was habituated to the reverse light cycle for 1 week and then underwent 5 weeks of fentanyl drinking in standard mouse cages. Cohort 4 then underwent 10-14 days of fentanyl abstinence prior to sacrifice for electrophysiological studies. This time point matches the period of abstinence the mice in cohort 1-3 experienced while undergoing fear conditioning.

### Drinking in the Dark (DiD) Model

On the first day of the drinking paradigm, mice were given either a bottle of plain water or fentanyl dissolved in drinking water beginning 3 hours into the dark cycle. The bottles were removed after 4 hours and replaced with drinking water until the next day. Mice underwent DiD for 5 days (Monday-Friday) with Saturdays and Sundays off. This cycle was repeated for 5 weeks, increasing the concentration of fentanyl in drinking water each week (10 ug/mL, 20 ug/mL, 30 ug/mL (for 2 weeks), then 40 ug/mL). Mice were housed in single-bottle cages, and the mice receiving fentanyl did not have access to normal drinking water during the 4-hour drinking sessions. Bottles were weighed at the beginning and end of each drinking session to determine the amount consumed. One bottle was filled with drinking water and set on an empty cage to account for drip.

Body weight was measured on the last day of each round of DiD.

### Opioid withdrawal behaviors

Opioid withdrawal behaviors were assessed as previously described ^25,26,34^. All animals were injected with 1 mg/kg naloxone following 4-5 hours of free access to 40 µg/mL fentanyl in water or water as a control. Immediately after injection, mice were placed in an open arena and withdrawal behaviors were assessed for 10 mins. The withdrawal behaviors evaluated included: escape jumps, paw tremors, jaw tremors, abnormal postures, wet dog shakes, grooming behaviors, and fecal boli number. Data were converted to a z-score for each behavior and a mean z-score of all behaviors together was calculated to generate a global withdrawal score for each mouse ^34,35^.

### Avoidance assays

The open field apparatus was a 40 cm x 40 cm clear plastic box with Plexiglass flooring in a sound attenuated chamber with a light source of 55 lux. Mice were placed in the center of the open field where locomotor activity was measured based on three zones: surround, center, and corners of the open field. The surround zone excluded the corners. Activity was analyzed with Omnitech Electronics Software (Omnitech Electronics Inc., Columbus, OH).

The elevated plus-maze (EPM) apparatus consisted of two open arms and two closed arms, all 77 cm x 77 cm and made of Plexiglass. All arms were connected to a central platform to form the apparatus, which stood 74 cm above the ground. Each mouse was placed on the central platform at the beginning of each trial, and an overhead camera recorded activity for 5 minutes. Time spent in each arm was analyzed with Ethovision software (Noldus, Netherlands).

The Light-Dark box was made of two compartments of equal size. The light compartment was illuminated to 300 lux. Mice were placed in the dark compartment, and an overhead camera recorded activity for 15 minutes. Videos were analyzed with Ethovision software (Noldus, Netherlands).

### Sucrose Preference Test (SPT)

SPT was performed according to published methods ^36^. Mice were moved to two-bottle choice cages for SPT and allowed to habituate for 5 days. 3 hours into the dark period, all mice were given one bottle of drinking water and one bottle of 1% sucrose in drinking water. The bottles were switched with one another 12 hours into the drinking period to account for side preference. The bottles were replaced with drinking water after 24 total hours. The same style of bottles were used for both fentanyl DiD and the SPT. The amount of sucrose consumed was measured by a sucrose preference ratio, dividing the volume of sucrose solution consumed by the total volume of fluid consumed.

### Fear Conditioning

Fear learning was conducted over a 4-day protocol. On day 1, mice were habituated to the fear conditioning chamber with a shock grid floor (Med Associates, Vermont, USA). On day 2, fear conditioning was performed with a 2 min baseline and 5 tone-shock pairings (tone 30s, 80 db, 3 kHz; shock 0.5 mA, 2 s). Mice then underwent 2 days of fear extinction in a novel context. Following a 2 min baseline, mice received 10 tone presentations (60 s, 80 db, 3 kHz). Behavior hardware was controlled by Ethovision XT and time spent freezing was calculated at baseline and during each tone presentation.

### Brain slice preparation

Brain slices were prepared for whole-cell electrophysiology as previously described ^37^. Briefly, mice were deeply anesthetized with isoflurane, decapitated, and brains were removed and placed into ice-cold sucrose aCSF [in mM: 194 sucrose, 20 NaCl, 4.4 KCl, 2 CaCl_2_, 1.2 NaH_2_PO_4_. 10 glucose, 26 NaHCO_3_] oxygenated with 95% O_2_ 5% CO_2_ for slicing. Brains were sliced coronally at 200-300 μm using a Leica VT1000 vibratome (Germany). Slices were then transferred to a holding container with oxygenated aCSF [in mM: 124 NaCl, 4.4 KCl, 2 CaCl_2_, 1.2 MgSO_4_. 1 NaH_2_PO_4_, 10 glucose, 26 NaHCO_3_] at 32 °C for at least 45 minutes. Slices were then transferred to a recording chamber and perfused with oxygenated aCSF [in mM: 124 NaCl, 4.4 KCl, 2 CaCl_2_, 1.2 MgSO_4_. 1 NaH_2_PO_4_, 10 glucose, 26 NaHCO_3_] at 30 °C at a constant rate of 2 mL/min.

### Whole-cell recordings

Recordings were conducted as previously described ^38,39^. Cells in which access resistance changed greater than 20% were excluded from analysis. 2-3 cells were recorded from each animal for each set of experiments. BLA principal neurons were identified by their anatomical location, low membrane resistance < 60 MΩ, and high capacitance > 100 pF. For measurements of action potential kinetics, the first resolvable, single evoked action potential was used to determine AP threshold, AP half-width, fAHP, and mAHP. Electrophysiology data was analyzed using Easy Electrophysiology (Easy Electrophysiology Ltd., London, UK).

### Drugs

Fentanyl citrate was obtained from Spectrum Pharmacy Products (New Brunswick, NJ, USA). Naloxone hydrochloride was obtained from Sigma-Aldrich (St. Louis, MO, USA).

### Statistics

All data were analyzed using Graph Pad Prism (version 10.0.02). For most studies males and females were analyzed separately due to significant differences in fentanyl consumption. Male and female data were collapsed for the electrophysiology studies because no sex differences were found in baseline properties. Data are reported as mean ± SEM. Comparisons were made using student’s t-test, regular or repeated measures 2-way ANOVA, or 3-way ANOVA depending on the number of variables. Šídák’s multiple comparisons test or Tukey’s multiple comparisons test were used for post-hoc analyses. Correlations were determined by calculating the Pearson Correlation Coefficient. Significance was defined as p ≤ 0.05.

## Results

### Fentanyl drinking

We developed a home-cage fentanyl drinking paradigm modeled on a single bottle DiD method. In our model, mice had access to increasing concentrations of fentanyl (10 – 40 µg/mL) dissolved in water over the course of 5 weeks for 4 hours. We chose to start at 10 µg/mL fentanyl because previous work has suggested that this concentration is not aversive to mice and rats, although less is known about higher concentrations ^11,40,41^. Control animals received a bottle of standard drinking water. Mice consumed increasing amounts of fentanyl by day over each week (**Figure 1A**, main effect of dose, F_6,192_ = 29.08, p < 0.0001). Notably, we observed sex differences with female mice consuming more fentanyl at the 30 µg/mL concentration where we saw peak consumption (**Figure 1B**, main effect of sex, F_1,145_ = 29.43, p < 0.0001, Šídák’s multiple comparison test, 30 µg/mL p = 0.021, p = 0.006, 40 µg/mL p=0.27). The overall volume of fluid consumed decreased in fentanyl consuming animals suggesting the taste of fentanyl may be slightly aversive at higher concentrations, particularly at highest concentration tested, 40 µg/mL, or alternatively that fentanyl reduces motivation to drink water at these concentrations (**Figure 1C & D**, time x treatment interaction, F_4,244_ = 7.83, p < 0.0001; main effect of treatment, F_1,61_ = 23.11, p < 0.0001). We also assessed changes in body weight over the course of the drinking paradigm. The body weight of all treatment groups increased over the course of the experiment and was unaltered by fentanyl consumption (**Figure 1E**, main effect of time, F_2.15,127_ = 102.0, p < 0.0001; main effect of treatment, F_1,59_ = 0.17, p = 0.67).

**Figure 1.**
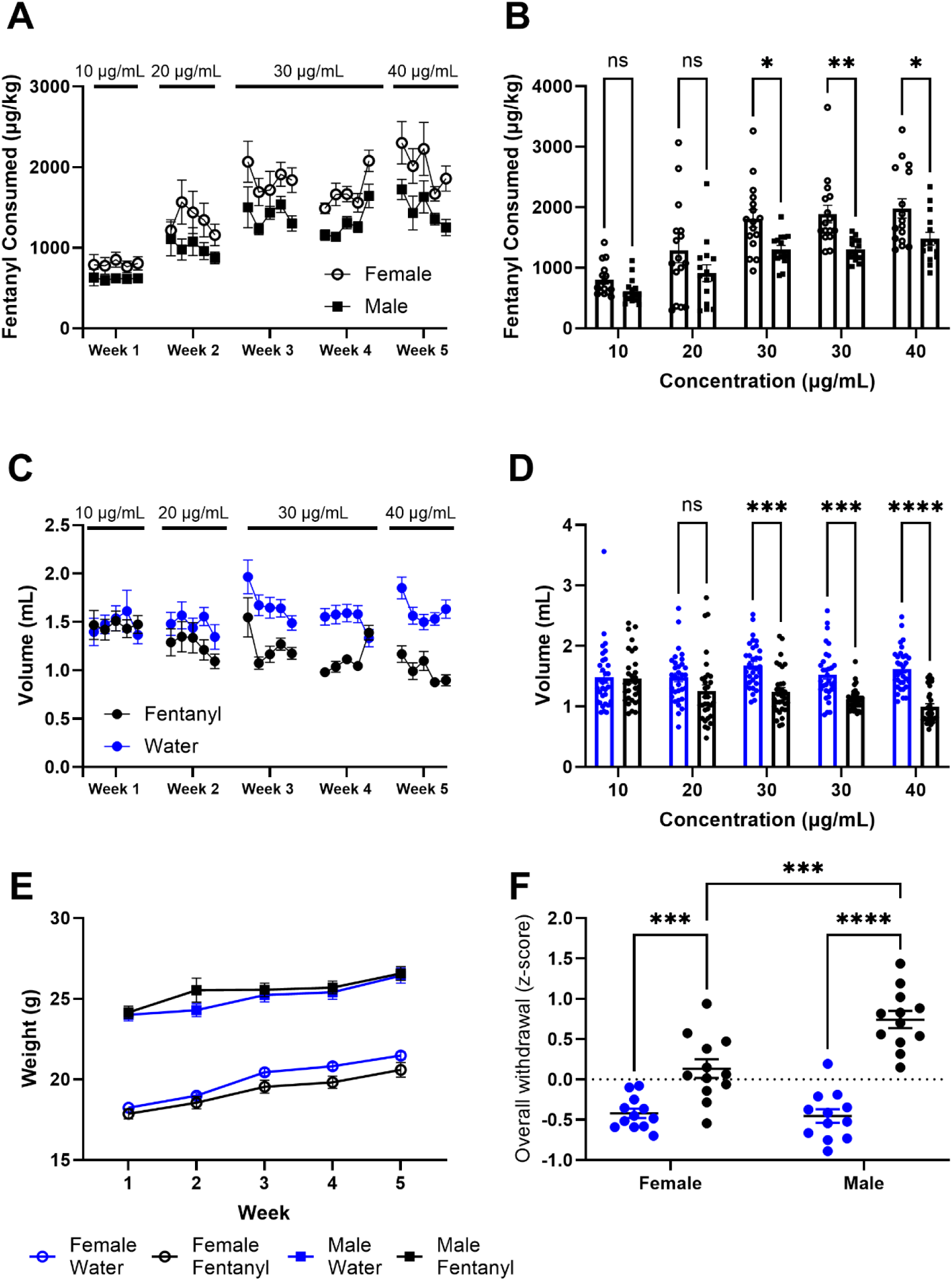
Volume and fentanyl consumed throughout 5-week fentanyl drinking paradigm and overall withdrawal behavior in female and male mice. **A.** Fentanyl consumed (µg/mL) by day in male and female mice. **B.** Average fentanyl consumed per week in male and female mice. Females consumed more fentanyl than males, especially at the 30 µg/mL dose. **C.** Volume consumed (mL) by day in fentanyl groups and water groups. **D.** Average volume consumed per week in fentanyl and water groups. Fentanyl groups consumed significantly less fluid at the 30 µg/mL dose and greater. **E.** Weight across 5-week paradigm with no differences observed between treatment groups. **F.** Overall withdrawal score (z-score) of fentanyl and water groups by sex. Fentanyl groups exhibited more withdrawal behaviors than water groups in both sexes. Males exhibited more withdrawal behaviors than females. Each point represents the mean ± standard error of the mean (SEM): * p<0.05, ** p<0.01, *** p<0.001, **** p<0.0001.

### Opioid withdrawal behaviors

To assess if mice consumed sufficient fentanyl to induce withdrawal signs and if there were sex differences in withdrawal symptoms following oral fentanyl consumption, we injected mice with 1 mg/kg naloxone immediately following the last 4 hour DiD session at the 40 µg/mL concentration of fentanyl. We assessed a variety of withdrawal behaviors including fecal boli production, escape jumps, paw tremors, wet dog shakes, abnormal posture, grooming, and jaw tremors. All behaviors were converted to z-scores to account for sex-specific expression of withdrawal behavior and allow for an unbiased assessment of overall withdrawal behaviors between male and female mice ^34^. To assess overall withdrawal severity, we calculated the mean z-scores for all withdrawal symptoms together. The fentanyl drinking group exhibited significantly more withdrawal behaviors than water drinking mice (**Figure 1F**, main effect of treatment, F_1,59_ = 121.7 p < 0.0001; Tukey’s multiple comparisons test, Males p < 0.0001, Females p = 0.0008). Males exhibited more withdrawal behaviors than females overall (sex x treatment interaction, F_1,59_ = 17.95, p < 0.0001; main effect of sex, F_1,59_ = 14.77, p =0.0003). See Supplement for an analysis of individual withdrawal behaviors (**Figure S1**).

### Fentanyl consumption alters avoidance behaviors

Data from our lab and other published studies have demonstrated protracted changes in avoidance behavior following opioid use in rodent models ^26,42–45^. One week after our five-week oral fentanyl administration paradigm, we assessed changes in avoidance behavior using 3 standard behavioral tests: open field (OF), elevated-plus maze (EPM), and light-dark box (LD). Given the baseline sex differences in fentanyl consumption, and our previous studies demonstrating sex differences in behavior in opioid withdrawal, we continued to analyze all behavior data stratified by sex ^26,27^. There were no significant differences in total distance traveled in the OF assay (**Figure 2A**, main effect of treatment, F_1,44_ = 0.30, p = 0.59). The female fentanyl group spent significantly less total time in the surround than the female water group (**Figure 2B**, main effect of treatment, F_1,44_ = 5.0, p=0.0292; Šídák’s multiple comparisons test, Male p=0.7901, Female p=0.0269). There were no differences between treatment groups in either males and females on time spent in the center (**Figure 2C**, main effect of treatment, F_1,44_ = 0.03, p = 0.86) or corners (**Figure 2D**, main effect of treatment, F_1,44_ = 1.86, p = 0.18). When examining the OF data across time, we found that female fentanyl mice spent less time in the surround than water mice, particularly at later time points (**Figure 2E** main effect of treatment, F_1,44_ = 5.08, p = 0.29). There were no differences in time spent in the center across the trial time (**Figure 2F**, main effect of treatment, F_1,44_ = 0.12, p = 0.73). However, the fentanyl mice spent significantly more time in the corner of the open field, particularly in the last 10 minutes of the trial (**Figure 2G**, treatment x time interaction effect, F_5,220_ = 4.01, p = 0.001). In the EPM test, we did not observe any significant differences in open arm entries, time in open arms, or latency to enter open arms in fentanyl or water controls (**Figure 2H, I, J, & K**). For the LD assay, we did not observe any treatment effects for latency to enter the light side (**Figure 2L**, main effect of treatment, F_1,44_ = 2.77, p=0.1035). The male fentanyl group spent less time on the light side of the box than the male water group, while all female mice spent a similar amount of time on the light side (**Figure 2M**, sex x treatment interaction, F_1,44_ =6.9, p=0.0118; Šídák’s multiple comparisons test, Male p=0.0468, Female p=0.3228). We did not observe treatment effects on the number of light side entries (**Figure 2N**, main effect of treatment, F_1,44_ = 0.12, p=0.7307).

**Figure 2.**
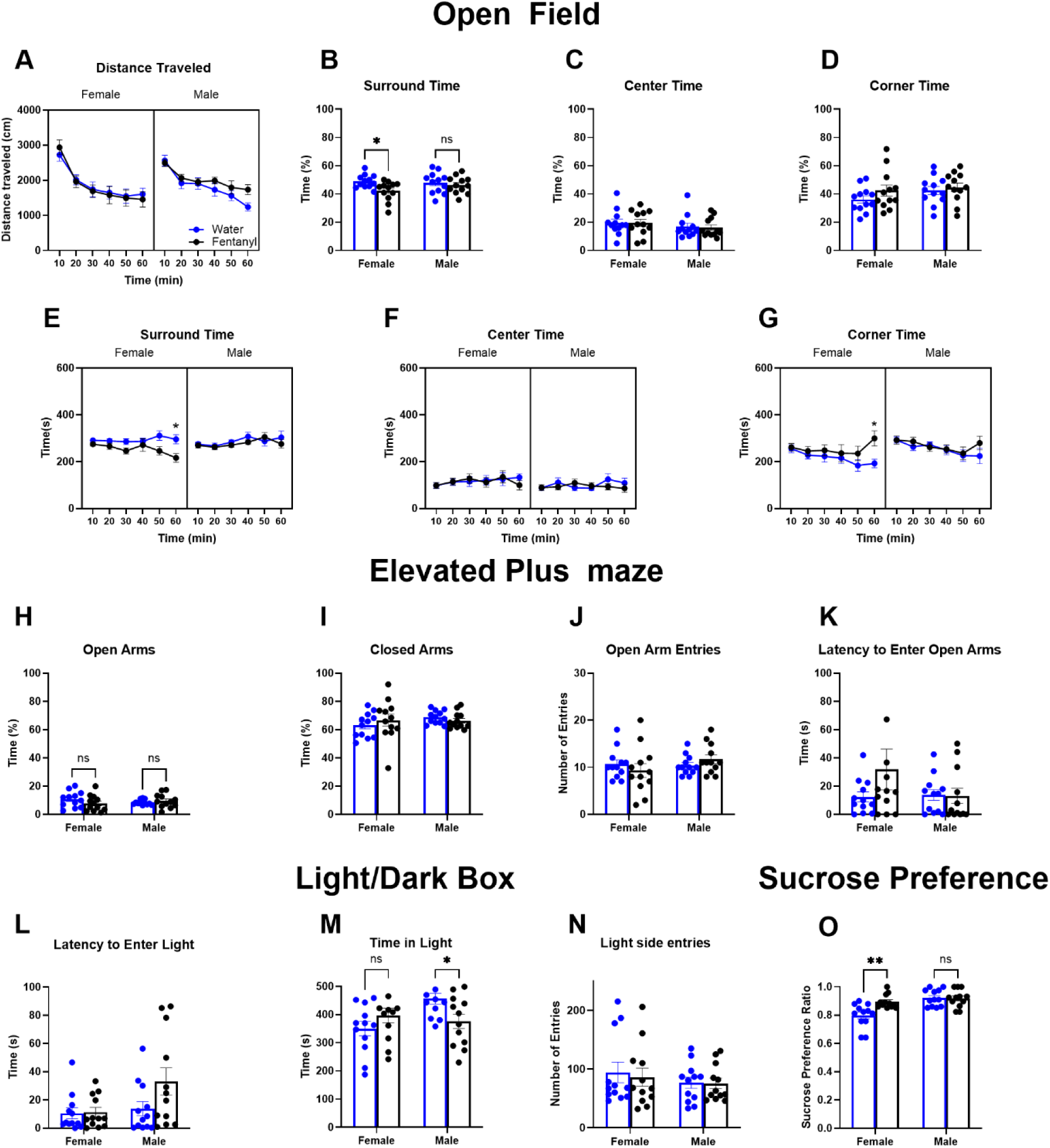
Behavioral assays following fentanyl drinking paradigm in male and female mice. **A, B, C, D, E, F, G.** Show behavior in the open field assay for locomotor activity. **A.** Treatment groups showed no differences in total distance traveled when binned in 10-minute increments across the 60-minute assay). **B.** Females showed decreased percent time spent in the surround in the fentanyl group, while males showed no differences across treatment groups. **C, D.** No differences were observed in the percent time spent in the center or corners between treatment groups for either sex. **E.** Fentanyl groups spent relatively less time in the surround region towards the end of the assay. **F.** Treatment groups showed no differences in percent time spent in center at any point throughout the assay. **G.** Fentanyl groups spent more time in the corner region at the end of the assay. **H, I, J, & K**. report behavior during the elevated-plus maze (EPM). **H & I.** There were no differences between sexes or treatments groups on time spent in either the closed or open arms of the EPM. **J**. There were no differences in open arm entries in the EPM. **K.** There were no differences in latency to enter the open arm in the EPM. **L, M, & N.** Show behavior in the light/dark box assay. **L.** There were no significant differences in the latency, in seconds, to enter the light side. **M.** The male fentanyl group spent less time in the light overall than the male water group, while females did not show significant differences. **N.** For both sexes, there were no differences in the number of times subjects entered the light side. **O.** Shows the sucrose preference test with 1% sucrose solution. The female fentanyl group showed a higher ratio of sucrose solution to total fluid consumed than the female water group, while males showed no differences. Each point represents the mean ± standard error of the mean (SEM): * p<0.05, ** p<0.01, *** p<0.001, **** p<0.0001.

### Fentanyl consumption alters motivation for reward in female mice

Substance use disorder can result in anhedonia or altered reward signaling ^46–48^. To assess changes in anhedonia or motivation for reward, we performed a sucrose preference test. Only female mice showed a difference in sucrose preference ratio based on treatment. Surprisingly, the female fentanyl group had a significantly greater sucrose preference than the female water group (**Figure 2O**, sex x treatment interaction, F_1,44_ =7.115, p=0.0107; main effect of sex, F_1,44_ =15.34, p=0.0003; main effect of treatment, F_1,44_ =6.181, p=0.0168; Šídák’s multiple comparisons test, Male p=0.9897, Female p=0.0014).

### Fentanyl consumption disrupts fear extinction

Previous studies have demonstrated that chronic opioid consumption alters normal fear learning and extinction ^28,29^. To determine if long-term oral fentanyl consumption disrupts fear learning, mice underwent a fear learning and fear extinction paradigm 10 days into abstinence from fentanyl. On day 1, mice were habituated to the fear conditioning chamber equipped with a shock floor. There were no baseline differences in distance traveled, average velocity, or time spent immobile (freezing) between the treatment groups (data not shown). On day 2, mice underwent fear learning. Total time spent freezing increased with each subsequent trial, but there were no significant differences between treatment groups in either male or female mice (**Figure 3A**, main effect of treatment, F_1,44_ = 0.50, p = 0.49). Mice next underwent fear extinction in a novel context, with a unique cage floor, walls, and scent cues. On day 1 of extinction, there were no differences in freezing time between the treatment groups (**Figure 3B**, main effect of treatment, F_1,44_ = 1.37, p = 0.25). However, on day 2 of the extinction trial, both male and female fentanyl drinking mice spent significantly more time freezing relative to the water controls (**Figure 3C**, treatment x tone presentation interaction effect, F_10.440_ = 3.01, p = 0.001, main effect of treatment, F_1,44_ = 15.6, p = 0.0003).

**Figure 3.**
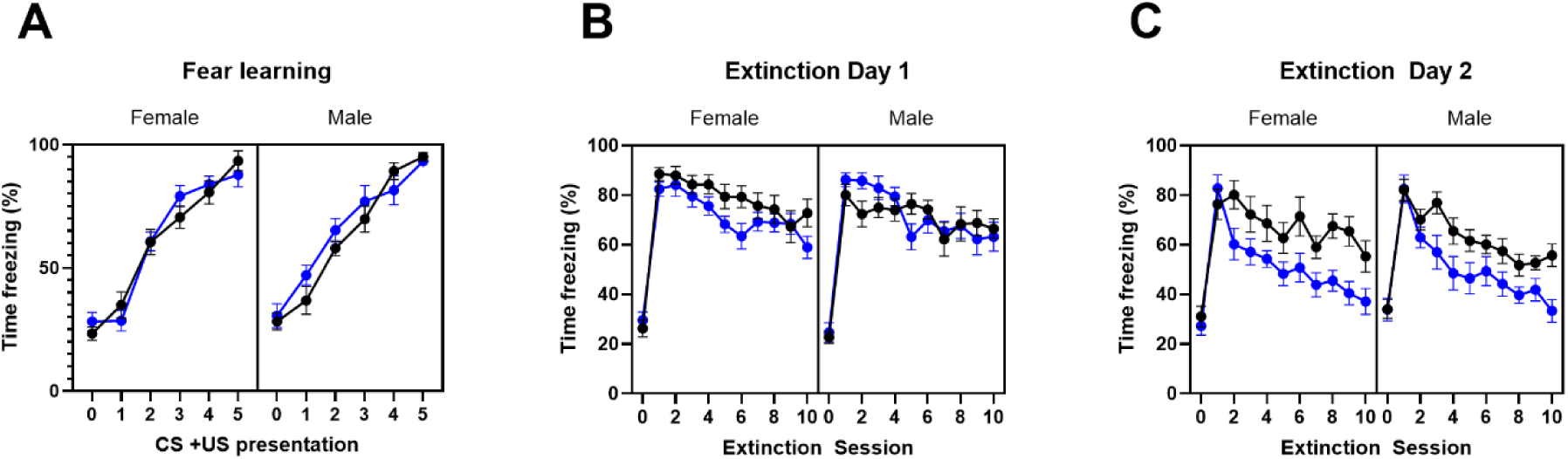
Fear extinction is impaired following fentanyl consumption. **A.** Percent time spent freezing during fear learning was consistent across treatment groups for both sexes. On extinction days, cues were played every two minutes for one minute. **M.** On the first day, there was no significant difference in freezing between each group. **N.** On the second day, the fentanyl groups for both sexes showed significantly more time freezing across all cues. CS + US presentation “0” and extinction session “0” denote baseline levels of freezing behavior. Each point represents the mean ± standard error of the mean (SEM): * p<0.05, ** p<0.01, *** p<0.001, **** p<0.0001.

**Figure 4.**
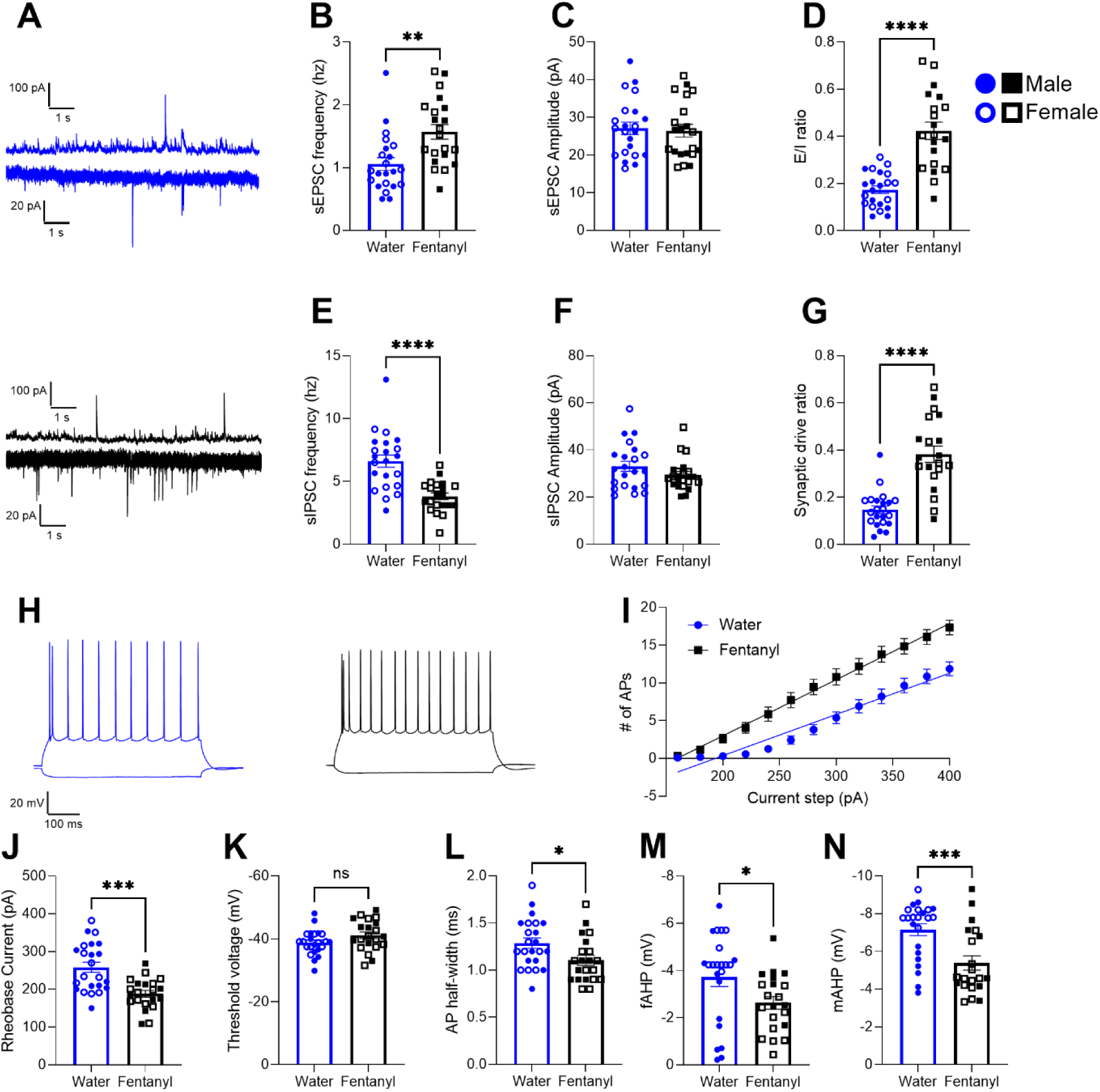
BLA neuron principal neuron physiology is altered following fentanyl self-administration. **A.** Representative sIPSCs traces (top) and sEPSCs traces (bottom) from water drinking (blue) and fentanyl drinking (black) mice. **B**. sEPSC frequency is increased following fentanyl administration. **C**. sEPSC amplitude is unchanged following fentanyl consumption. **D**. BLA principal neuron E/I ratio is increased following fentanyl consumption. **E**. sIPSC frequency is decreased following fentanyl consumption. **F**. sIPSC frequency is unchanged following fentanyl consumption. **G**. Synaptic drive ratio is increased following fentanyl consumption. **H**. Representative voltage traces following hyperpolarizing (−100 pA) and depolarizing (400 pA) current injection. **I**. BLA principal neurons are more excitable to depolarizing current injection following fentanyl consumption. **J**. Rheobase current injection is reduced following fentanyl consumption. **K**. Action potential threshold voltage is unchanged following fentanyl consumption. AP half-width (**L**), fast after hyperpolarization potential (**M**), and medium after hyperpolarization potential (**N**) are reduced following fentanyl consumption. Open symbols represent females and closed symbols represent males. Each point represents the mean ± standard error of the mean (SEM): * p<0.05, ** p<0.01, *** p<0.001, **** p<0.0001. (n=7 mice, 2-3 cells per animal for both current clamp and voltage clamp experiments)

### Fentanyl consumption enhances BLA principal neuron excitability and excitatory balance

Previous studies have found that increased BLA principal neuron excitability contributes to fear extinction deficits ^49,50^. Given our observations that fentanyl consumption disrupted normal fear extinction learning, we next investigated BLA principal neuron physiology. All BLA recordings occurred at least 10-14 days into fentanyl abstinence in a separate set of animals that did not undergo behavioral testing. Membrane properties, including membrane resistance, access resistance, and capacitance were similar between all groups (**Table** 1). We first assessed excitatory/inhibitory (E/I) balance in voltage clamp. We did not observe any significant sex differences in this data set, so male and female data were combined. There was a significant increase in sEPSC frequency in the fentanyl group (**Figure 5A & B**, (t)=3.35, p = 0.002) but no change in sEPSC amplitude between groups (**Figure 5C**, (t)=0.26, p = 0.79). We observed a decrease in sIPSC frequency in the fentanyl group (**Figure 5A & E**, (t)=4.92, p < 0.0001), with no change in sIPSC amplitude (**Figure 5F**, (t)=1.41, p = 0.17). Due to the opposing changes in sIPSC and sEPSC frequency observed in the fentanyl group, we found an increased E/I ratio in the fentanyl group (**Figure 5D**, (t)=6.44, p < 0.0001). To further assess the relative strength of excitatory and inhibitory inputs, we calculated the synaptic drive ratio for each cell. The synaptic drive ratio is calculated as the frequency of sEPSCs multiplied by the amplitude of sEPSC divided by the frequency of sIPSCs multiplied by the amplitude of sIPSCs. We observed a significant increase in the synaptic drive ratio of the fentanyl group (**Figure 5G**, (t)=6.32, p < 0.0001). In a sperate experiment, we performed the same voltage-clamp recordings in mice after they underwent the fear conditioning paradigm and after 1 week of re-exposure to fentanyl and found many of the same changes in E/I balance and changes in excitatory and inhibitory input (**Figure S3 A-F**).

**Figure 5.**
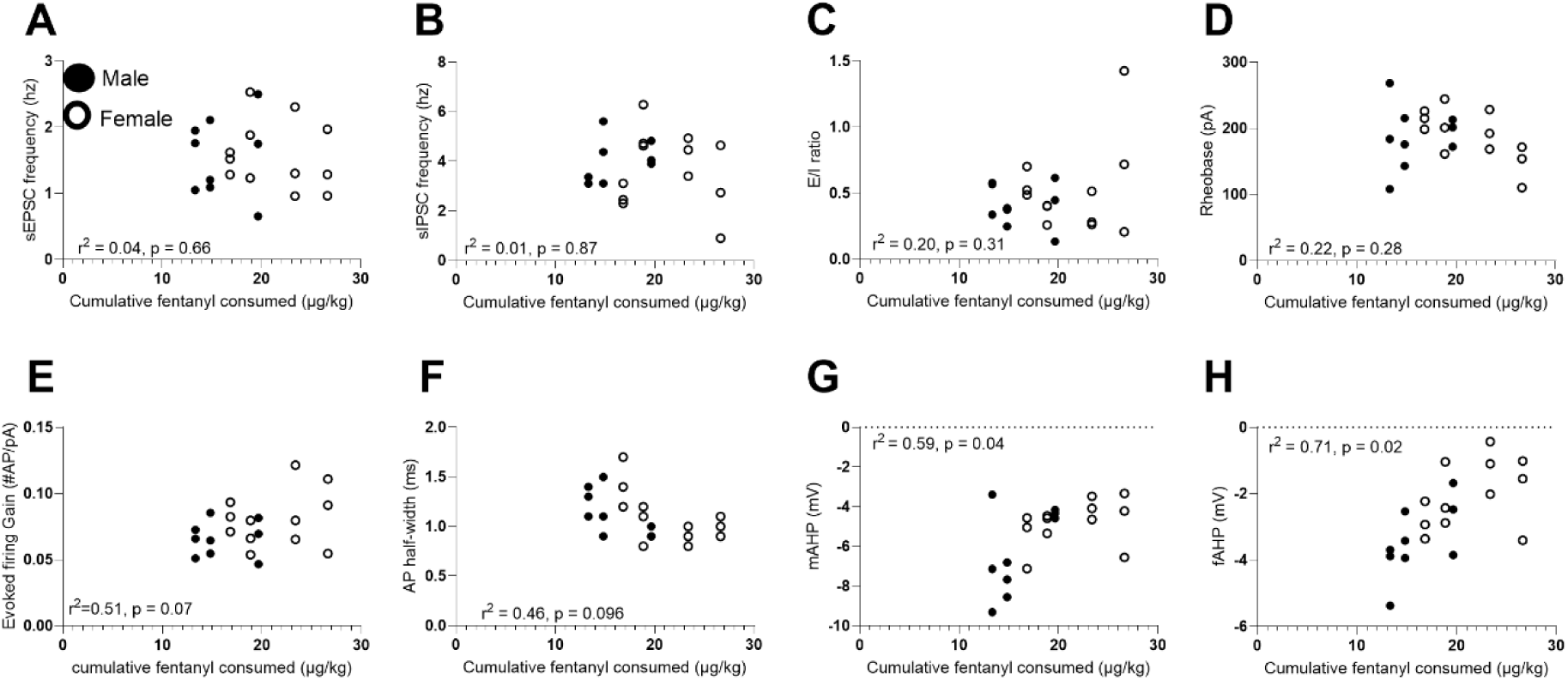
Correlations between cumulative fentanyl consumed and electrophysiological properties for each animal. sEPSC frequency (**A**), sIPSC frequency (**B**), E/I ratio (**C**), and rheobase (**D**) are poorly correlated with cumulative fentanyl consumed. There is a trend towards a correlation between evoked AP firing gain (**E**) and AP half-width (**F**) and cumulative fentanyl consumed. Both mAHP (**G**) and fAHP (**H**) are well correlated with fentanyl consumed. (n = 7 mice, 2-3 cells per animal)

We next assessed the excitability of BLA principal neurons in current clamp. While holding neurons at −75 mV, we found that the BLA principal neurons from the fentanyl group had a significantly reduced rheobase current (**Figure 5J**, (t)=4.30, p < 0.0001). We also tested neuronal responses to hyperpolarizing and depolarizing current injection (**Figure 5H**). BLA neurons from the fentanyl group had a significantly increased number of evoked action potentials per depolarizing current step (**Figure 5J**, current step x fentanyl interaction effect, F_12,504_ = 8.89, p < 0.0001). We also assessed the kinetics of evoked action potentials. Both fentanyl and control groups had similar AP threshold potentials (**Figure 5K**, (t)=1.73, p = 0.09). AP half-width was decreased following fentanyl consumption (**Figure 5L**, (t)=2.38, p = 0.022). We also found that both fAHP (**Figure 5M**, (t)=2.34, p = 0.031) and mAHP (**Figure 5N**, (t)=3.58, p = 0.001) were reduced following fentanyl consumption. These data suggest that both intrinsic changes in ion channel conductance on BLA principal neurons and extrinsic changes in excitatory and inhibitory connectivity may underlie increased excitability of these cells following fentanyl consumption. In a sperate experiment, we performed the same voltage-clamp recordings in mice after they underwent the fear conditioning paradigm and after 1 week of re-exposure to fentanyl and found many of the same changes rheobase current and evoked action potential firing (**Figure S3 G-H**).

### Fentanyl consumption correlates with changes in BLA AP kinetics

We next assessed whether any of changes in electrophysiological properties correlated with an individual animal’s cumulative fentanyl consumption. We chose to correlate changes with cumulative fentanyl consumption to reflect the entire history of fentanyl exposure rather than a single week when fentanyl consumption varies depending on fentanyl concentration. We did not observe significant correlations between cumulative fentanyl consumption and sEPSC frequency (**Figure 5A**), sIPSC frequency (**Figure 5B**), E/I ratio (**Figure 5C**), or rheobase current (**Figure 5D**). There was a trend towards a correlation between cumulative fentanyl consumption and evoked AP firing gain (**Figure 5E**) and AP half-width (**Figure 5F**). However, we did find correlations between cumulative fentanyl consumed and both mAHP (**Figure 5G**) and fAHP (**Figure 5H).** We did not find any significant correlations between any of our electrophysiological measurements and withdrawal severity (**Figure S2**).

## Discussion

### Oral fentanyl consumption model

Most studies of opioid consumption have focused on intravenous administration, although other routes of administration, such as vaping or oral ingestion, are increasingly more common for fentanyl ^6,7,21,22,24^. We adapted a DiD model to begin to study the consequences of chronic oral fentanyl consumption. We found that mice consumed larger amounts of fentanyl with increasing concentrations of fentanyl from 10-40 µg/mL. This was despite an overall decrease in the total volume of fluid consumed with increasing fentanyl concentrations. Fentanyl consumption stabilized between the 30 and 40 µg/mL concentration which may suggest that mice titrate their consumption to achieve a desired effect, which was previously proposed in an oral oxycodone self-administration paradigm ^12,51^. Alternatively, fentanyl may have an aversive taste, reduce motivation to drink fluid, or induce greater psychomotor effects at higher concentrations, which may reduce consumption. Notably, we did not observe sedative effects of fentanyl, which suggests that consumption is not limited by sedation at these doses (data not shown). We observed significant sex differences in fentanyl consumption, with females consuming more fentanyl than males at the 30 µg/mL concentration. However, we have observed that female mice consume more liquid (g/kg) than male mice in general ^52,53^. While studies investigating sex differences in oral fentanyl consumption are extremely limited, our work agrees with existing opioid self-administration literature that consistently shows increased opioid self-administration in females versus males ^11,16,54–56^.

While all mice in the fentanyl group exhibited similar withdrawal symptoms after naloxone-precipitated withdrawal, we found that male mice showed significantly more withdrawal signs. This is surprising because female mice consumed more fentanyl, which suggests that withdrawal severity is not solely dependent on the amount of fentanyl consumed. Our findings agree with recent work from our laboratory that found that males experienced more severe withdrawal from fentanyl, albeit in a model of non-contingent fentanyl administration. Interestingly, this same study demonstrated that the kappa-opioid system may play a larger role in somatic withdrawal in females vs. males ^35^. While clinical studies generally report that women experience worse withdrawal symptoms than men, it is important to note that these studies are typically limited to heroin or illicit use of prescription opioids rather than fentanyl ^57,58^. Further, sex as a biological variable explores the range of potential biological variability vs. explicit differences stratified to the sexes ^59,60^.

### Fentanyl consumption alters avoidance behaviors, reward-seeking, and fear extinction

While the physical symptoms of withdrawal, such as vomiting, diarrhea, and chills, subside shortly after the cessation of opioid consumption, protracted withdrawal symptoms, such as anxiety and depression, can last for months after the cessation of opioid use ^61,62^. Our lab has previously demonstrated protracted withdrawal symptoms in mice 6 weeks after naloxone-precipitated morphine withdrawal ^26^. In the current study, we found sex-dependent differences in avoidance behavior after fentanyl administration. Female mice in the fentanyl group spent more time in the corner in the last 10 minutes of the open field assay, suggesting impaired habituation. We found that male mice that consumed fentanyl spent significantly less time in the light-side of the light/dark box and had a trend toward a longer latency to enter the light-side, while females were unchanged. This suggests that fentanyl-consumption may increase anxiety-like behavior, albeit in an assay and sex-specific manner. Multiple clinical studies have established a connection between anxiety disorders and opioid use disorder, and the association between anxiety and opioid use disorder is especially strong in women ^63–65^. Our data suggests that fentanyl use itself may increase anxiety-like behaviors, although future studies will need to examine more protracted timepoints several weeks after the cessation of opioid use.

Anhedonia is a common feature of opioid use disorder and often serves as a strong predictor of relapse ^66,67^. We assessed anhedonia using the sucrose-preference test and found increased sucrose preference in the female fentanyl-consuming group, but no change in males. This contrasts with previous work on the effects of opioids on anhedonia, and suggests our fentanyl model enhances motivation for reward, at least in females. This may be due to the fast clearance of fentanyl from the blood relative to morphine, which may differentially alter GABAergic transmission in the VTA ^68^. Alternatively, our results may reflect a ceiling effect due to the high preference for the 1% sucrose solution used in our study, and lower concentrations of sucrose may produce different results.

In humans, OUD is associated with mood disorders and PTSD ^69,70^. However, the causal relationship between OUD and PTSD is unclear. Conditioned fear extinction is commonly used to model the associative aspects of PTSD in preclinical animal studies and previous work has demonstrated reduced cue-induced extinction learning following chronic morphine exposure and that morphine pretreatment enhanced associative fear learning ^28,29^. We found that oral fentanyl exposure did not impact the acquisition of fear learning. However, fentanyl exposure impaired cued extinction learning on the second day of extinction learning, demonstrating that chronic fentanyl consumption disrupts long-term extinction memory. These effects were observed 10 days after the last fentanyl exposure, so they do not reflect the effects of active opioid exposure, and all fentanyl would be cleared by this time point ^68^.

### Fentanyl consumption alters BLA plasticity

The basolateral amygdala is one brain region that is critical for cued fear extinction and reward learning ^49,71,72^. We found that BLA principal neurons had greater excitatory inputs and were more excitable following fentanyl consumption. Importantly, previous work has demonstrated that increased BLA principal neuron excitability contributes to fear extinction deficits ^73,74^. While it is unclear which specific inputs to BLA principal neurons were altered, we observed both a decrease in inhibitory inputs and an increase in excitatory inputs. µ-ORs are robustly expressed in GABAergic lateral paracapsular neurons, and µ-OR activation inhibits these cells ^75^. Chronic fentanyl consumption may disrupt these GABAergic projections to BLA principal neurons to impair fear extinction ^76^. µ-ORs are also expressed in the PFC, which is a major excitatory input to the BLA. While acute µ-OR activation decreases excitatory inputs from the PFC to BLA, chronic fentanyl exposure may alter PFC plasticity to increase excitatory inputs to the BLA ^77,78^. A recent study investigating changes in BLA plasticity following chronic intermittent ethanol exposure (CIE) and withdrawal found that CIE induced similar increases in excitability in BLA principal neurons projecting to both the BNST and NAcc ^79^. While we did not investigate specific projection populations in the BLA, together these findings may suggest that BLA hyperexcitability is a common mechanism of drug exposure and/or withdrawal.

The increased excitability we observed in BLA principal neurons may be due to intrinsic factors in addition to the extrinsic changes in excitatory inhibitory balance. We observed decreased AP half-width, fAHP, and mAHP following fentanyl consumption and withdrawal. In fact, mAHP and fAHP were strongly correlated with the cumulative amount of fentanyl consumed by each mouse, which may suggest these factors are directly related to fentanyl consumption rather than withdrawal severity. Faster AHPs and repolarization promote faster firing rates and may explain the observed increased excitability in this study ^80^. While we did not examine individual currents that underlie these changes, it is likely there are changes in BK and SK currents along with changes in Ca^2+^ buffering and voltage-gated calcium channel expression ^81^. While changes in BLA excitability and AP kinetics have not been studied in the context of opioids to our knowledge, a number of studies have found reduced mAHP and fAHP amplitude in lateral amygdala and BLA principal neurons following acute stressors and this change was mediated by reduced expression of BK channels in these neurons ^82–84^. A similar mechanism may underlie our observations following fentanyl consumption. While these findings may be important to understanding how fentanyl exposure affects retention of aversive memories, future studies are needed to further investigate molecular and circuit mechanisms underlying these changes.

It is important to note a few limitations of this model and study. First, it is not possible to investigate motivation to consume fentanyl in this model. While mice can choose the amount of fentanyl to drink in our DiD paradigm, animals did not have access to normal drinking water. Future studies could use a 2-bottle choice model to assess preference for fentanyl over normal drinking water. However, the current model allows for the study of consequences of non-contingent fentanyl consumption and avoids the technical pitfalls of intravenous self-administration in mice. Second, this model studied multiple aspects of opioid use, including fentanyl consumption, spontaneous withdrawal, precipitated withdrawal, and abstinence which makes it difficult to determine if the observed physiological and behavioral changes are due to fentanyl consumption itself, withdrawal, abstinence, or a combination of all factors. While future studies could investigate withdrawal or consumption alone, it is important to note that people with opioid use disorder often go through cycles of use, withdrawal, and abstinence ^85^. While the fentanyl drinking paradigm used in this study may complicate interpretation, it is also likely more translatable to the clinical course of OUD. Third, the effects of fentanyl and withdrawal on avoidance behaviors were relatively modest and differed based on assay and sex. Previous studies in our lab have found stronger effects on avoidance behavior following opioid withdrawal ^26^. This may reflect more modest daily fentanyl intake in our model or the relatively short time the mice spent undergoing precipitated withdrawal. While the effects of our fentanyl paradigm on avoidance behaviors were relatively modest, we found stronger effects of fear extinction. This may suggest that our fentanyl model produces stronger effects on neurocircuitry mediating responses to traumatic stressors rather than baseline anxiety levels.

In summary, we developed a mouse model of chronic voluntary home-cage oral fentanyl consumption that results in disruptions to anxiety-like behaviors, reward motivation, fear extinction, and BLA physiology all of which have been implicated in OUD in humans. Future studies are needed to investigate how fentanyl disrupts key brain circuitry involved in these processes to help develop new treatments for OUD.

**Table 1.**
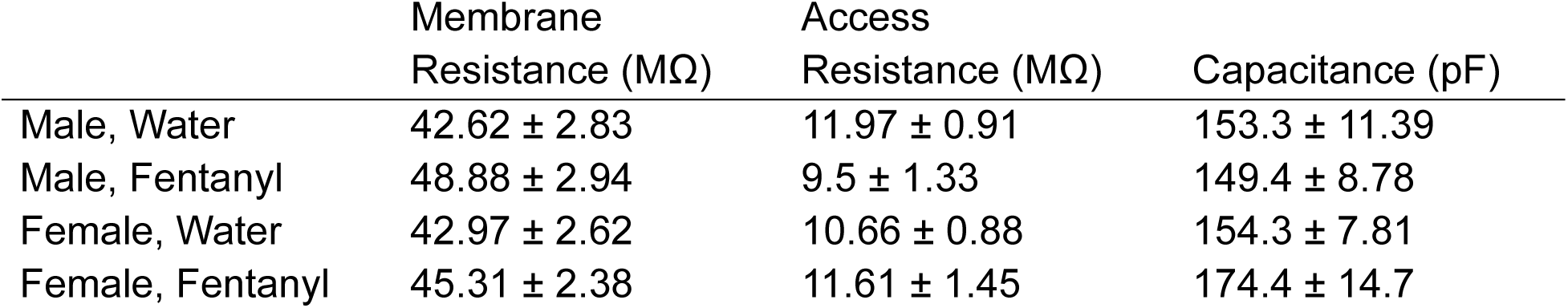
Membrane properties of BLA principal neurons. Data is expressed as mean ± standard error of the mean (SEM).

## Author Contributions

A.M.D designed and performed experiments, analyzed data, and wrote the manuscript.

G.K designed and performed experiments, analyzed data, and wrote the manuscript.

Z.A.M. designed experiments, wrote the manuscript, and provided funding for the study.

All authors approve the final manuscript and agree to be accountable for all aspects of the work.

## Supplemental Data

**Figure S1.**
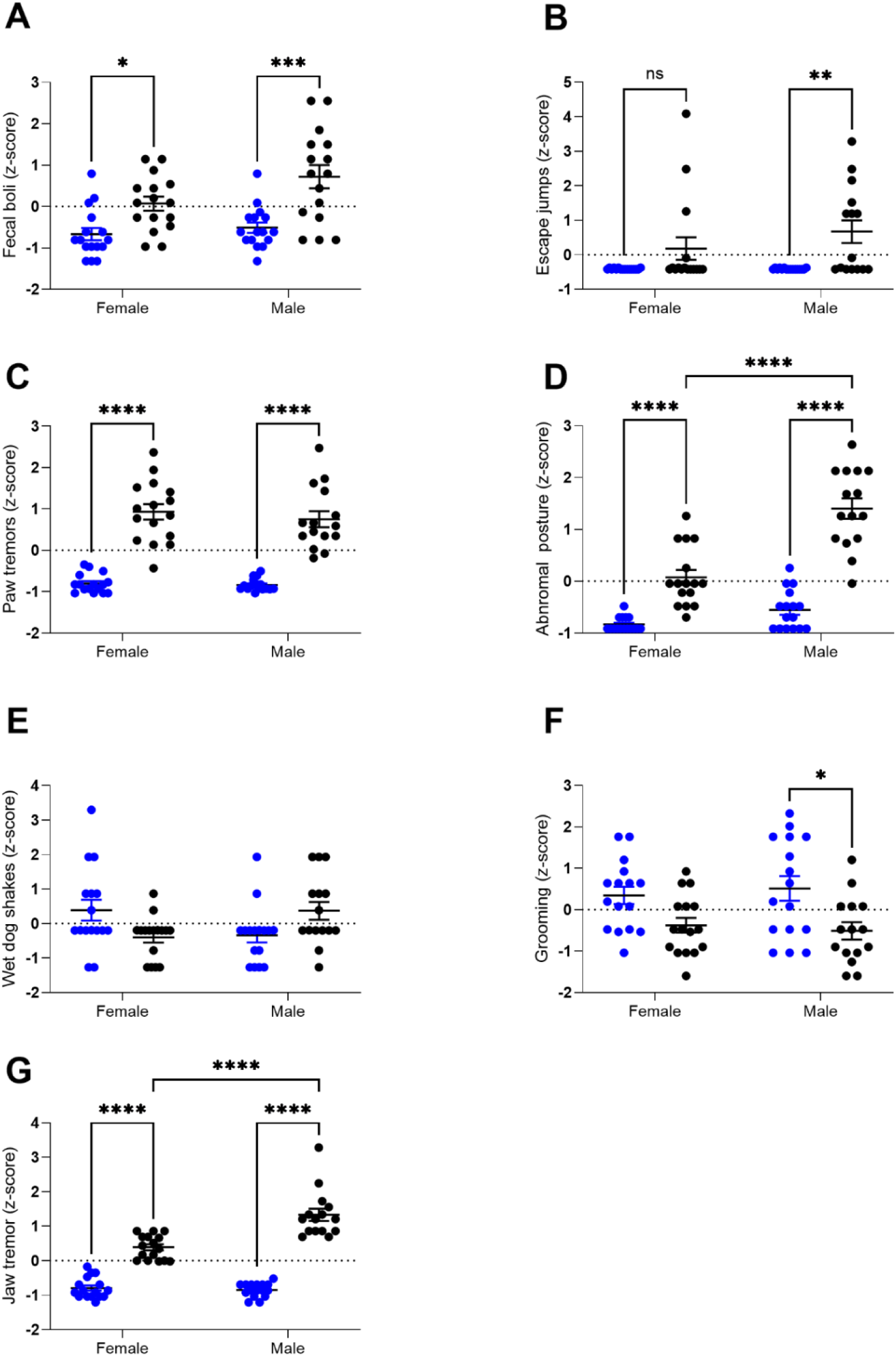
Long-term fentanyl consumption shows sex differences in withdrawal behaviors. **A** Male and female fentanyl group produced more fecal boli than water groups for both sexes. We observe a significant effect of sex driven by male mice (main effect of sex, F_1,60_ = 4.49, p = 0.04). **B** Only males exhibited more escape jumps in the fentanyl group. Females showed no differences between treatment groups. **C** Both male and female fentanyl groups had increased paw tremors than water groups. **D** Both male and female fentanyl groups had more instances of abnormal posture than water groups, but male fentanyl group had more instances than female fentanyl group (main effect of sex, F_1,60_ = 39.01, p < 0.0001). **E** We did not observe changes in wet dog shakes due to fentanyl treatment. However, we observed a significant interaction effect. (treatment x sex interaction, F_1,60_ = 9.96, p = 0.003) **F** Grooming behavior was significantly reduced following fentanyl treatment, although the effect was greater in males than females. **G** Both male and female fentanyl groups had increased jaw tremors, but male fentanyl group revealed more jaw tremors than female fentanyl group (main effect of sex, F_1,60_ = 17.46, 17.46, p < 0.0001). Error bars represent the mean ± standard error of the mean (SEM): * p<0.05, ** p<0.01, *** p<0.001, **** p<0.0001.

**Figure S2.**
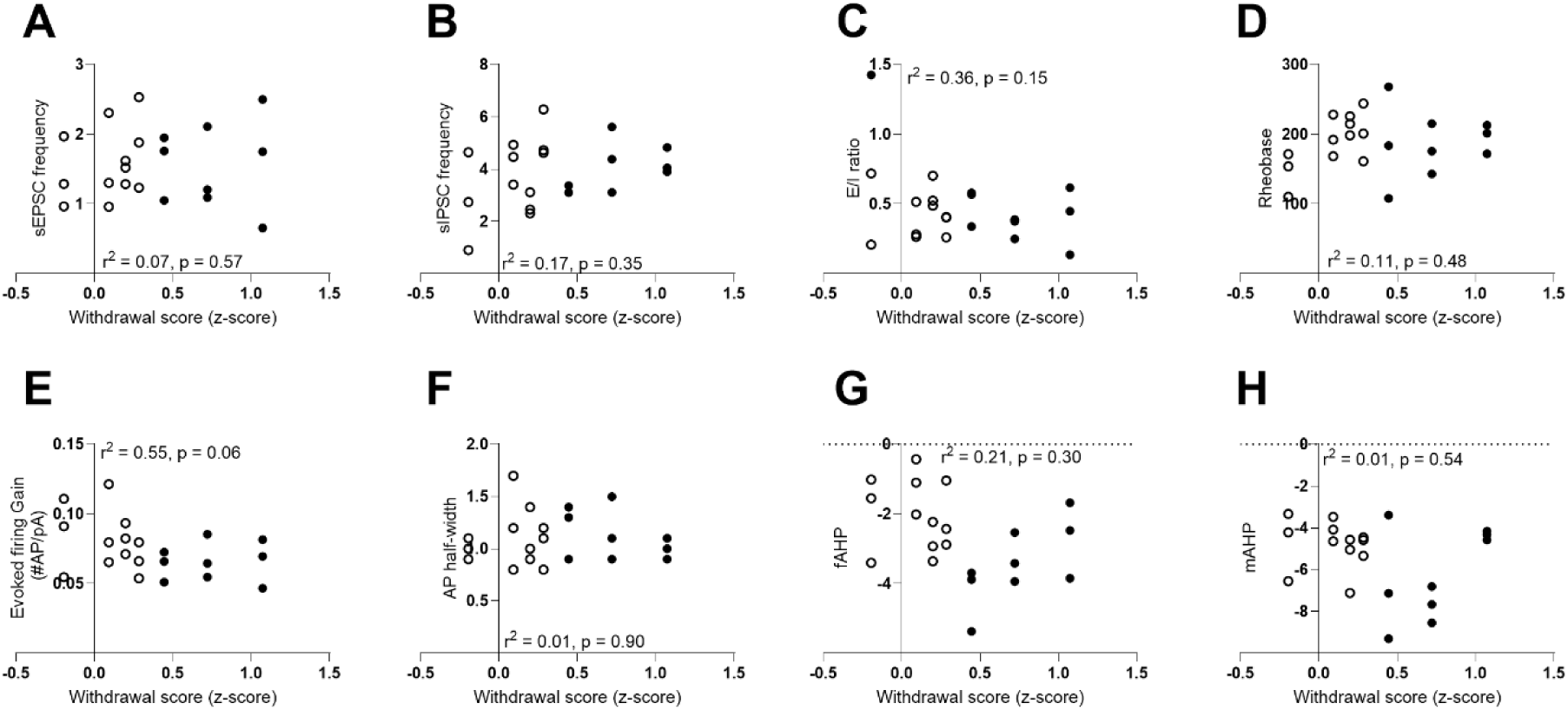
Correlations between average precipitated withdrawal score severity and electrophysiological properties. We did not observe correlations between overall withdrawal severity and sEPSC frequency **(A)**, sIPSC frequency **(B)**, E/I ratio **(C)**, Rheobase **(D)**, Evoked firing gain **(E)**, AP half-width **(F)**, fAHP **(G)**, mAHP **(H)**.

**Figure S3.**
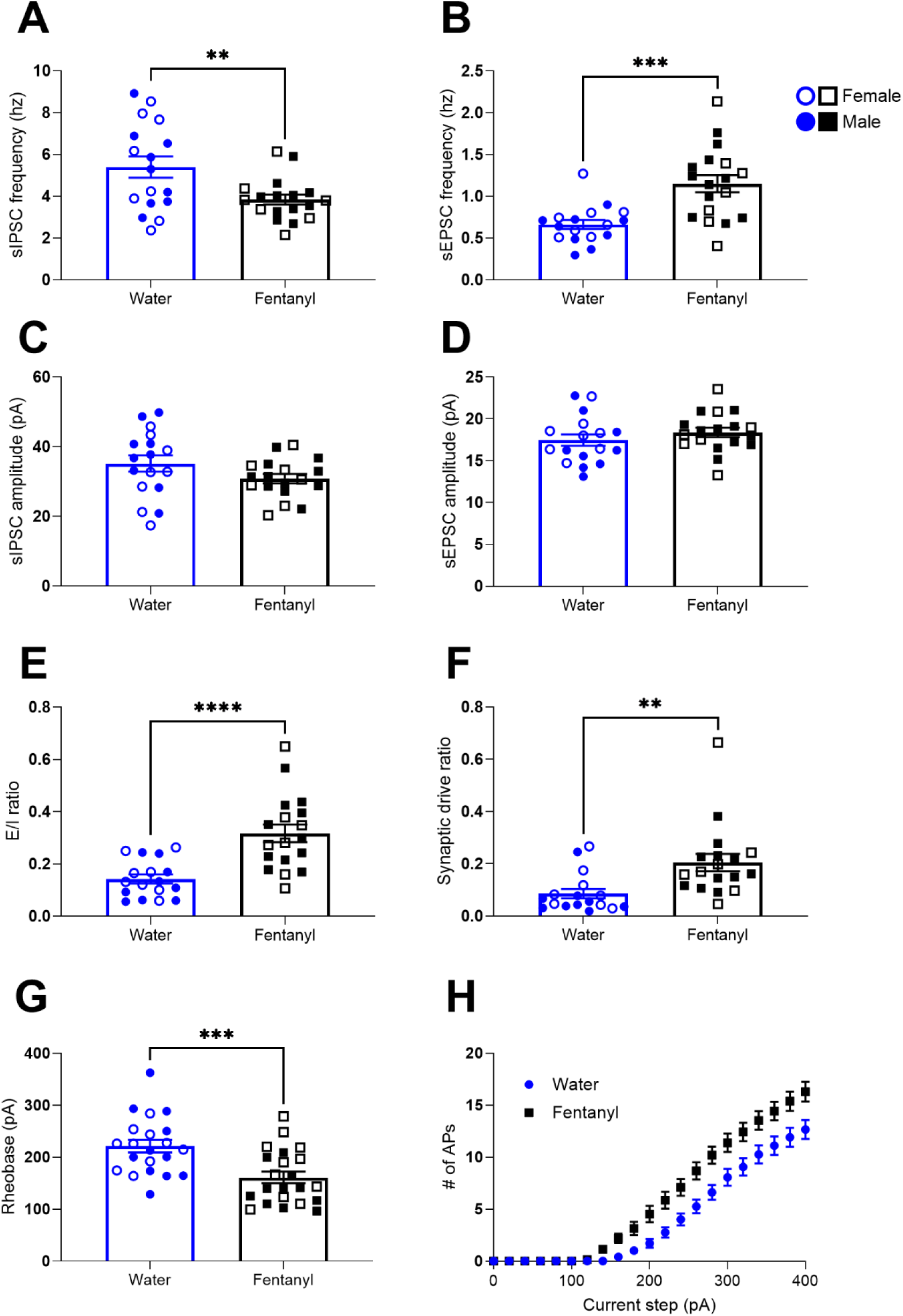
Excitatory and inhibitory inputs are altered following fear conditioning and 1 week of fentanyl reinstatement. For these experiment mice were recorded 3-5 days into abstinence. **A** sIPSC frequency is reduced in fentanyl mice. **B** sEPSC frequency is increased in fentanyl drinking mice. Neither sIPSC amplitude **(C)** or sEPSC amplitude **(D)** are altered by fentanyl consumption. **E** E/I ratio and synaptic drive ratio **(F)** are increased following fentanyl consumption. **G** Rheobase current is reduced following fentanyl consumption. **F** Evoked action potential firing is increased in fentanyl mice with increasing current injection. Closed symbols represent males and open symbols represent female data. Data is expressed as mean ± SEM. * p<0.05, ** p<0.01, *** p<0.001, **** p<0.0001

